# Associative white matter tracts selectively predict sensorimotor learning

**DOI:** 10.1101/2023.01.10.523345

**Authors:** S. Vinci-Booher, D.J. McDonald, E. Berquist, F. Pestilli

**Affiliations:** Department of Psychological and Brain Sciences, Program for Neuroscience, Indiana University, Bloomington, Indiana, United States; Department of Psychology and Human Development, Vanderbilt University, Nashville, Tennessee, United States; Department of Statistics, University of British Columbia, Vancouver, British Columbia, Canada; Department of Psychology, Center for Perceptual Systems, Center for Theoretical and Computational Neuroscience, Center for Aging Populations Sciences, Center for Learning and Memory, University of Texas at Austin, Austin, Texas, United States

## Abstract

Human learning is a complex phenomenon that varies greatly among individuals and is related to the microstructure of major white matter tracts in several learning domains, yet the impact of the existing myelination of white matter tracts on future learning outcomes remains unclear. We employed a machine-learning model selection framework to evaluate whether existing microstructure might predict individual differences in the potential for learning a sensorimotor task, and further, if the mapping between the microstructure of major white matter tracts and learning was selective for learning outcomes. We used diffusion tractography to measure the mean fractional anisotropy (FA) of white matter tracts in 60 adult participants who then underwent training and subsequent testing to evaluate learning. During training, participants practiced drawing a set of 40 novel symbols repeatedly using a digital writing tablet. We measured drawing learning as the slope of draw duration over the practice session and visual recognition learning as the performance accuracy in an old/new 2-AFC recognition task. Results demonstrated that the microstructure of major white matter tracts selectively predicted learning outcomes, with left hemisphere pArc and SLF 3 tracts predicting drawing learning and the left hemisphere MDLFspl predicting visual recognition learning. These results were replicated in a repeat, held-out data set and supported with complementary analyses. Overall, results suggest that individual differences in the microstructure of human white matter tracts may be selectively related to future learning outcomes and open avenues of inquiry concerning the impact of existing tract myelination in the potential for learning.

**Significance statement:** A selective mapping between tract microstructure and future learning has been demonstrated in the murine model and, to our knowledge, has not yet been demonstrated in humans. We employed a data-driven approach that identified only two tracts, the two most posterior segments of the arcuate fasciculus in the left hemisphere, to predict learning a sensorimotor task (drawing symbols) and this prediction model did not transfer to other learning outcomes (visual symbol recognition). Results suggest that individual differences in learning may be selectively related to the tissue properties of major white matter tracts in the human brain.

---

The structural architecture of the human brain contains large bundles of myelinated fibers called white matter tracts. The white matter tracts carry communications among cortical regions and functional components of the brain. Evidence from animal models suggest that these white matter bundles may be selectively related to human learning, such that the myelination of some bundles may be more related to learning a particular task than that of other bundles ^1–3^. Seminal work in a murine model demonstrated that myelination of major white matter tracts was necessary for behavioral changes, i.e., learning, following controlled stimulation of cortical activity ^2^. Similarly, in the murine model, myelination of major white matter tracts was necessary for learning that followed controlled behavioral training experiences ^1,3^. Whether myelination followed cortical stimulation or behavioral training, myelination was necessary for learning and selective, meaning that it was promoted in some tracts but not others.

As of today, selective myelination of white matter tracts in humans as a result of learning a particular task has been elusive. On the contrary, changes in white matter across multiple tracts is most often reported after training on a task, such as juggling or reading ^4–11^. For example, one significant study measured tract microstructure in children before and after an intensive reading intervention ^11^. Prior work from a variety of methodologies, including correlational, post-mortem anatomy, and case studies, suggested that the intensive reading intervention would promote microstructure changes in two specific white matter tracts: the Arcuate fasciculus (Arc) and the Inferior Longitudinal Fasciculus (ILF) ^12–14^. Instead, the study found non-specific, widespread changes across multiple tracts. Currently, the human literature comparing pre- and post-intervention white matter has not detected a tract-selective relationship between white matter and learning.

A different approach to testing tract selectivity is investigating the relationship between current tract myelination and future learning outcomes. This approach exploits the existing myelination, which occurred over the course of the lifespan, instead of shorter-term experimental interventions. Studies taking this approach have demonstrated that the microstructure of white matter tracts can be used to predict future learning, such as semantic learning ^15,16^, foreign language learning ^17^, auditory tone learning ^18^, visuomotor adaptation ^19^, and face-name learning ^20^. In each case, studies suggest that a relatively small, select group of tracts predicts learning, i.e., tract-selectivity, and that the tracts most predictive of learning one task may not also be most predictive of learning a second task, i.e., task-selectivity. Related work suggests that connectivity predicts learning sensorimotor tasks, such as learning to play novel piano sequences ^21,22^. Critically, studies testing the relationship between existing tract myelination and future learning outcomes have focussed on tracts of interest (investigating only one or a few tracts) and have often tested a single learning outcome (investigating a single task without testing transfer of learning across behaviors). As a result, current evidence for tract- and task-selectivity in humans remains inconclusive.

Here, we tested tract-selectivity in humans learning a sensorimotor task using machine-learning and a model-selection framework. Instead of implementing a training protocol and measuring changes in white matter as a result of learning, we used the myelination that occurred over the course of the lifespan to predict future learning. We tested tract-selectivity by investigating the relationship between the microstructure of major white matter tracts (**Figure 1a**) and individual differences in performance on two learning outcomes following a single sensorimotor training task (**Figure 1b**). Two learning outcomes were included so as to allow us to test the different task-selectivity of white matter tracts to different learning outcomes.

**Figure 1.**
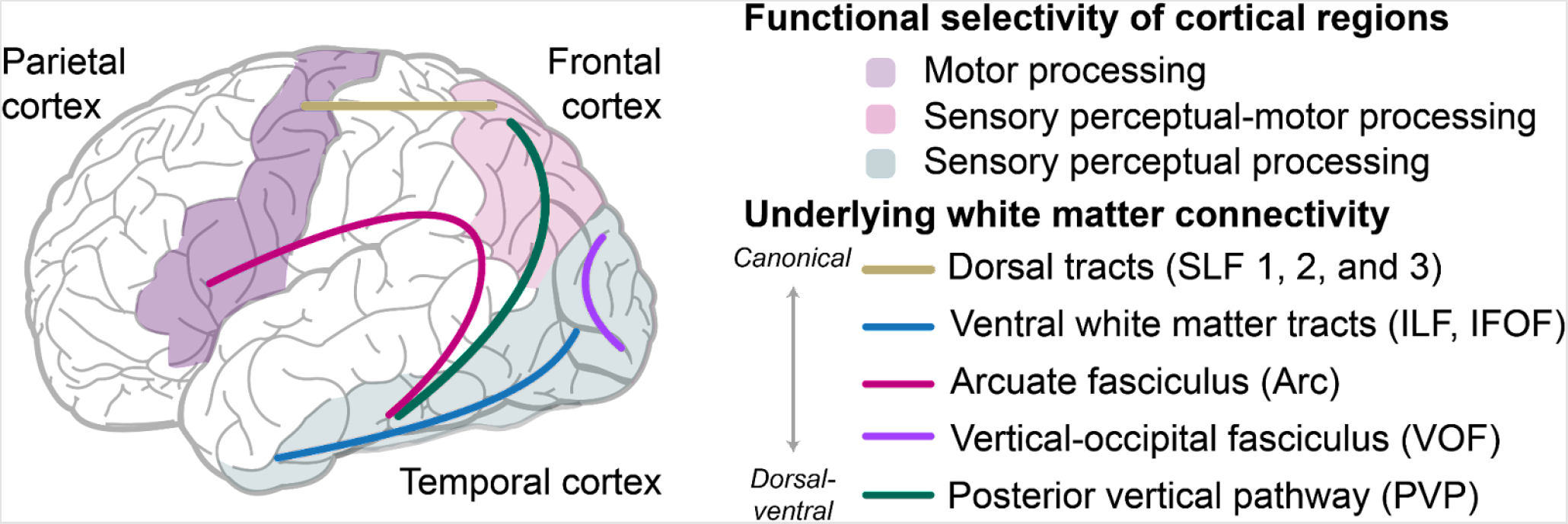
Background and hypotheses. Major white matter tracts connect frontal, parietal, and temporal cortical regions. Prior works have demonstrated that frontal activation is associated with motor movements during drawing symbols, temporal activation is associated with perceptual processing of the symbols produced, and parietal activation is associated with both motor and perceptual processing ^30^. The hypothesis of the current work is that a select group of these major white matter tracts will predict individual differences in learning to draw novel symbols and not visual recognition learning, and vice versa. **SLF:** superior longitudinal fasciculus; **PVP:** posterior vertical pathway; **Arc:** arcuate fasciculus; **ILF:** inferior longitudinal fasciculus; **IFOF**: inferior fronto-occipital fasciculus; **MDLFspl:** middle longitudinal fasciculus connection to the superior parietal lobe; **MDLFang:** middle longitudinal fasciculus connection to the angular gyrus; **TPC:** temporal to parietal connection; **pArc:** posterior arcuate fasciculus.

## Results

We investigated the mapping between the microstructure of white matter tracts and two learning outcomes that arise from the same training experience to test both tract- and task-selectivity. We measured white matter microstructure in 60 adult participants who then completed a sensorimotor training task that required them to practice drawing a set of 40 novel symbols repeatedly using a digital writing tablet. After the training, participants were tested on their ability to visually recognize the 40 symbols (**Figure 2a**). We estimated drawing learning by measuring the draw duration of each symbol drawing trial and calculating the slope of draw duration over the practice session (**Figure 2b**). We estimated visual recognition learning by measuring performance accuracy in an old/new 2-AFC visual recognition task. Accuracy was selected over reaction time because both metrics demonstrated learning and accuracy captured more individual variability than reaction time (see **Supplementary Information**).

**Figure 2.**
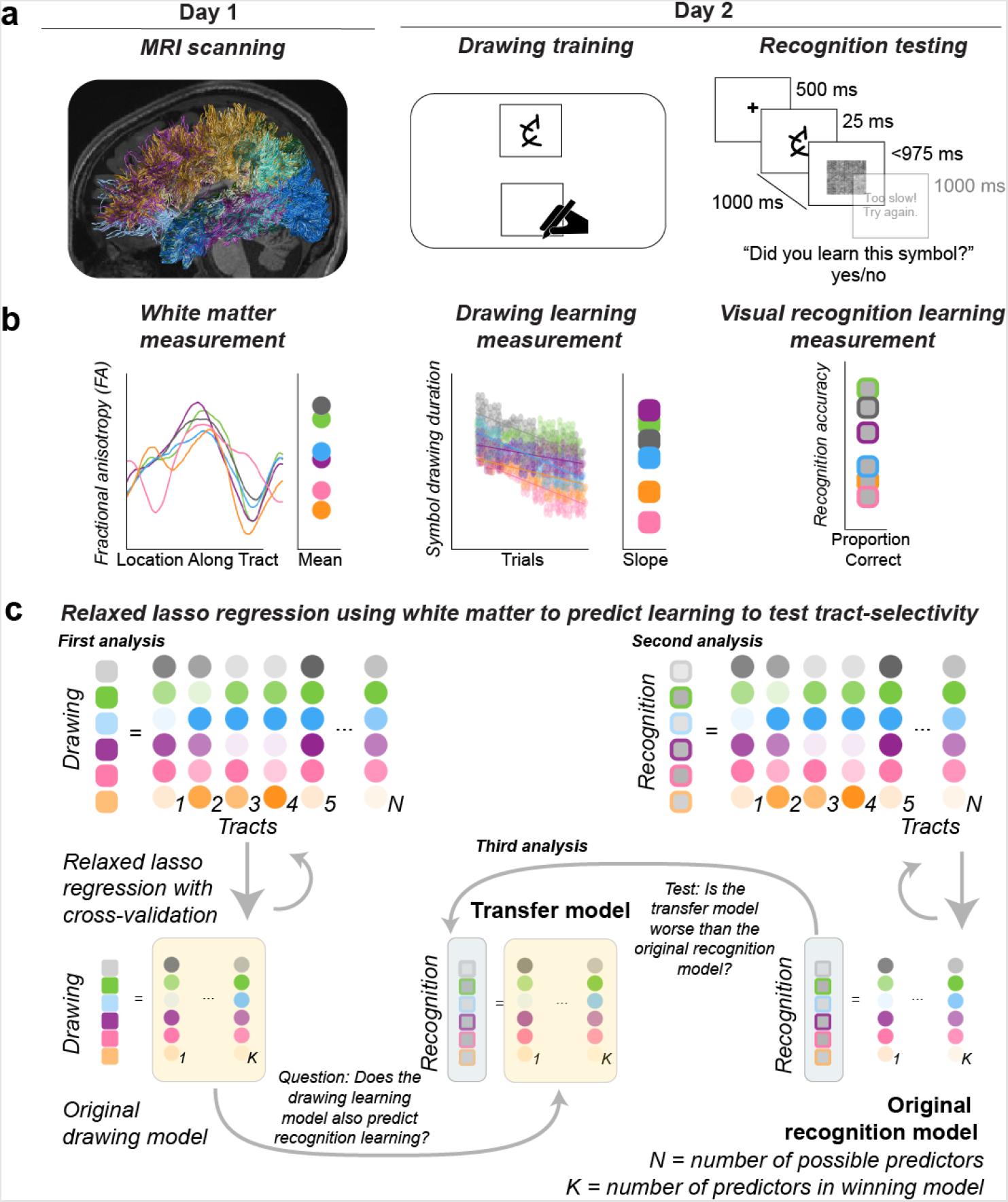
Experimental procedure, measurements, and modeling approach. **a**. Overall procedure. All participants completed an MRI session before completing a session of drawing training and recognition testing. On Day 1, diffusion MRI data were collected at the in order to perform diffusion tractography and estimate tissue microstructure. On Day 2, participants completed 30 minutes of drawing training, including 40 novel symbols each drawn 10 times in random order (drawing training) followed by an old/new 2-AFC visual recognition test (recognition testing). **b**. Calculations of white matter and learning measurements. First, a measurement of mean fractional anisotropy (FA) was obtained for each tract using a tractprofiles approach and averaging across the tract profile. Second, a measurement of drawing learning was obtained by estimating the linear slope of draw duration over trials. Third, a measurement of visual recognition learning was obtained by estimating the proportion of correct responses on the visual recognition test. The white matter measurements on day 1 were then used to predict the measurements of drawing learning and visual recognition learning on day 2. **c**. Modeling approach. A relaxed lasso regression was used to identify the group of tracts that was most predictive of individual differences in drawing learning and, separately, most predictive of visual recognition learning. After predictors were selected for drawing learning and for visual recognition learning separately, we tested to see if the drawing learning model transferred to visual recognition learning (Transfer model). We found that the Transfer model was worse at predicting visual recognition learning than the model selected for visual recognition learning (Original model), demonstrating task-selectivity because the model selected for learning to draw symbols did not also predict learning to visual recognize symbols.

We employed relaxed-lasso (RL) methods ^31,32^ to specify a model to test the hypothesis that a selective group of white matter tracts predicts (in cross-validation terms) learning to draw novel symbols (**Figure 2c**). The RL method allows testing models where the best set of parameters (the tracts’ Fractional Anisotropy, or FA) is selected to best fit the data. Critically, RL does not guarantee identification of a small subset of tracts, but it is capable of selecting all tracts, if that provides the best fitting model ^32^.

Furthermore, we used RL to test that a different group of tracts predicted learning to visually recognize the symbols. We conducted one RL regression to predict drawing learning and a second to predict visual recognition learning as an initial assessment of task-selectivity (see our third analysis for an additional test of task-selectivity) from the microstructure of white matter tracts.

We directly tested a set of 22 white matter tracts that connect cortical regions known to support motor and sensory processing during symbol drawing ^30,33–40^ (**Figure 1a**). The Superior Longitudinal Fasciculus (SLF 1 and 2 combined, SLF 3) directly connects frontal and parietal cortices where neural processing is largely associated with motor planning and control, respectively ^23^. Within the ventral cortex where neural processing is largely associated with sensory perception, the Inferior Longitudinal Fasciculus (ILF) directly connects occipital and temporal cortices ^24,25^, the Inferior Fronto-Occipital Fasciculus (IFOF) connects occipital and prefrontal cortices ^24,26,27^. Between the dorsal motor and ventral perceptual cortices, the arcuate fasciculus (Arc) and the posterior vertical pathway (PVP) directly connect the ventral perceptual cortex with the dorsal motor cortex. The Arc directly connects temporal and frontal cortices ^28,29^; the PVP directly connects temporal and parietal cortices and might be better thought of as a collection of four tracts: the Posterior Arcuate (pArc), the Temporal-to-Parietal Connection to the Superior Longitudinal Fasciculus (TPC), the Middle Longitudinal Fasciculus Connection to the Angular Gyrus (MDLFang), and the Middle Longitudinal Fasciculus Connection to the Superior Parietal Lobe (MDLFspl) ^30^. We also included two additional vertical tracts, the Vertical-Occipital Fasciculus (VOF) in the posterior cortex and the Frontal Aslant Tract (FAT) in the anterior cortex. The left and right hemispheres were kept separate for each of these 11 tracts, for a total of 22 white matter tracts tested.

We tested our hypothesis that a select group of white matter tracts would predict drawing learning in three ways (**Figure 2c**). In the **first analysis**, we sought to demonstrate that the microstructure of some tracts was more predictive of learning than other tracts, i.e., tract-selectivity. To do this, we specified an initial model that we called the original drawing learning model using a RL regression to select a set of white matter tracts that explained the most variance in drawing learning. In the **second analysis**, we constructed a second model that we called the original recognition learning model using a second RL regression to select tracts that explained the most variance in visual recognition learning, more so than other tracts, i.e., tract-selectivity. In the **third analysis**, we sought to directly test that the tracts identified for drawing learning were selective for drawing, i.e., task-selectivity. To do this, we evaluated whether the relationship between pre-training white matter microstructure and drawing learning would translate to a second learning outcome, namely visual recognition learning. All analyses were also conducted with a more extended set of tracts using an exploratory approach, however we found results consistent with the theory-driven selection of 22 tracts. We report the results of the theory-driven analysis in the main text and provide the results of the exploratory approach in **Supplementary Information**. Additionally, we report a complementary analysis using simple (marginal) linear regressions in **Supplementary Information** to evaluate the ability of the microstructure of any tract to independently predict each learning outcome.

### First analysis: Predicting drawing learning from the microstructure of major white matter tracts

We used a RL and model-selection via cross-validation to identify the subset of white matter tracts that best predicted sensorimotor learning, i.e., drawing learning. We hypothesized that the microstructure of a select group of white matter tracts prior to learning would predict learning to draw novel symbols. The results of our first analyses demonstrated tract-selectivity: the left pArc and left SLF3 selectively predicted learning to draw novel symbols while the left MDLFspl selectively predicted learning to visually recognize those symbols. The RL analysis selected the tracts that, together, explained the most variance in drawing learning using a leave-one-out cross-validation model selection procedure ^31,32^. We included 22 potential predictors in the lasso regression, including the microstructure of each PVP tract (pArc, TPC, MDLF-ang, MDLF-spl) as well as tracts within the dorsal cortex (SLF 1 and 2 combined as one tract, SLF 3) and tracts within ventral cortex (ILF, IFOF). We also included three additional vertical tracts, one in the posterior cortex (VOF) and one in the anterior cortex (FAT) and the arcuate fasciculus (Arc), for a total of 22 potential predictors that correspond to 22 white matter tracts (11 left and 11 right hemisphere). Results of the RL analysis supported our hypothesis: individual differences in the microstructure of PVP white matter tracts predicted drawing learning, specifically the left pArc but also the left SLF 3 (**Table 1; Figure 3a**). The microstructure of both tracts positively predicted drawing learning, and this result was replicated in a held-out dataset (**Table 1**) and in complementary marginal linear regressions (see **Supplementary Information**).

**Table 1.** Models selected for each learning outcome using relaxed lasso regression. Models were selected using RL. Summary statistics are reported for the selected models estimated via OLS using the final model predictors.

| Response Variable | Predictor | $\beta$ | S.E. | $R^2$ | Adj. $R^2$ |
| --- | --- | --- | --- | --- | --- |
| Drawing learning | Left pArc | 0.2118 | 0.2654 | 0.1180 | 0.0788 |
|  | Left SLF3 | 0.1772 | 0.2600 | - | - |
| Drawing learning (repeat dataset) | Left pArc | 0.1968 | 0.2517 | 0.0969 | 0.0568 |
|  | Left SLF3 | 0.1448 | 0.2569 | - | - |
| Visual recognition learning | Left MDLFspl | -0.6290 | 1.4424 | 0.0662 | 0.0247 |
|  | Left TPC | -0.4025 | 1.1971 | - | - |
| Visual recognition learning (repeat dataset) | Left MDLFspl | -0.7562 | 1.4981 | 0.0416 | 0.0208 |

**Figure 3.**
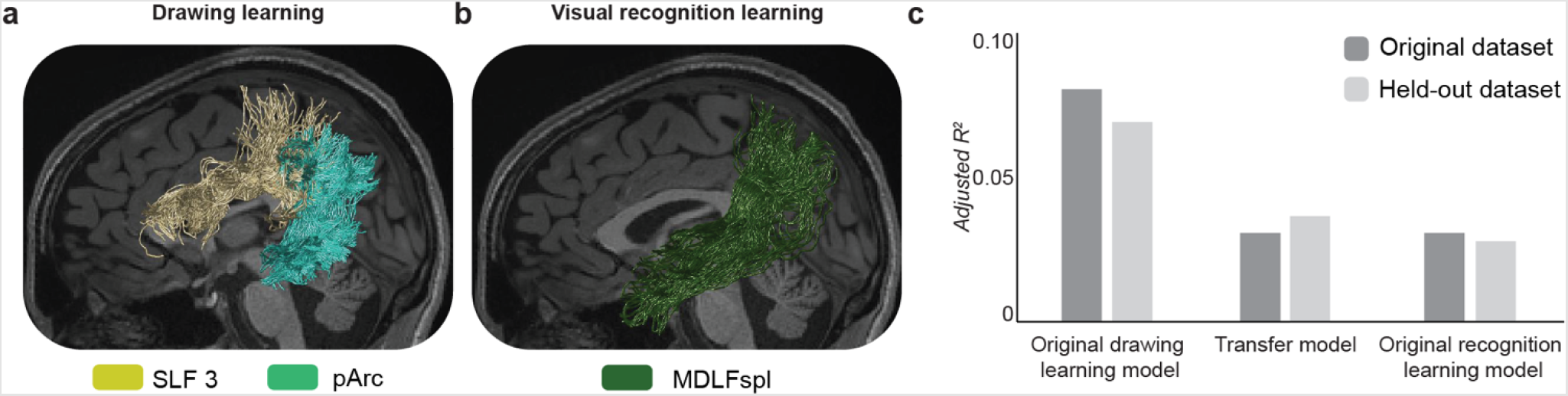
Associative white matter tracts selectively predict learning.**a**. Drawing learning. The left pArc and left SLF 3 selectively predicted drawing learning. A relaxed lasso (RL) regression analysis selected the left pArc and left SLF 3 from a set of 22 potential tracts as the tracts that explained the most variance in drawing learning after training using a leave-one-out cross-validation model selection procedure **b**. Visual recognition learning. A second RL regression revealed that the left MDLFspl predicted visual recognition learning. **c**. Task-selectivity. A transfer model was constructed that used the tracts selected for predicting drawing learning (a), the left pArc and left SLF 3, to predict visual recognition learning. A Cox test and a J-test both confirmed that the model selected for drawing learning did not transfer to predicting a second learning outcome, i.e.., visual recognition learning. The Cox test demonstrated that the original recognition learning model was a better fit than the transfer model to predict visual recognition learning. The J-test demonstrated that combining the transfer and original recognition models to predict visual recognition learning resulted in a model fit that was not significantly better than the original recognition model. **SLF 3:** third segment of the superior longitudinal fasciculus; **pArc:** posterior arcuate; **MDLFspl:** middle longitudinal fasciculus connection to the superior parietal lobe.

The RL regression optimized to predict drawing learning revealed that two left hemisphere tracts selectively predicted drawing learning: the left pArc and left SLF 3 (**Table 1; Figure 3a**). These two tracts were selected from 22 potential tracts from multiple competing models ^32^, suggesting tract-selectivity. With drawing learning as the dependent variable, the winning model included only the left pArc and the left SLF3, with a cross-validated model R^2^ = 0.4799. This result was replicated in a held-out, repeat dataset: left pArc and the left SLF 3, with a cross-validated model *R*^*2*^ = 0.4582. The relationships between the microstructure of the left pArc and left SLF3 were positive, such that participants with higher FA in those tracts were participants who were the quickest at learning to draw the novel symbols.

### Second analysis: Predicting visual recognition learning from the microstructure of major white matter tracts

A RL regression was also performed to predict visual recognition learning from the microstructure of pre-training white matter tracts (**Figure 2b**). Based on evidence from functional neuroimaging that indicated that the distribution of function across cortex during visual perception of known symbols was similar to the distribution of function while drawing symbols ^33,43–45^, our hypothesis for the white matter tracts that would significantly predict visual recognition was the same as our hypothesis for the tracts that would significantly predict drawing learning: we expected that tracts within the posterior vertical pathway would predict visual recognition learning.

An RL model was specified that predicted visual recognition learning (accuracy) given the 22 predictors previously used to select the model for drawing learning. Results of the RL analysis supported our hypothesis: individual differences in the microstructure of PVP white matter tracts predicted visual recognition learning, specifically the left MDLFspl (**Figure 3b**). Using the original dataset, two left hemisphere tracts were selected with a cross-validated model to predict visual recognition learning: left MDLFspl and left TPC (*R*^*2*^ = 0.4601) (**Table 1**). This result was partially replicated in the held-out dataset; the same analysis applied to the repeat dataset identified only the left MDLFspl, with a cross-validated model *R*^*2*^ = 0.3511. The results of the simple (marginal) linear regressions did not reveal a significant relationship between the microstructure of the left MDLFspl and visual recognition learning for any individual tract (see **Supplementary Information**). The only tract identified in both the original and repeat dataset was the left MDLFspl. In both analyses the relationship was negative, such that participants with higher FA in the left MDLFspl were participants who were the slowest at learning to draw the novel symbols.

### Third analysis: Directly testing the task-selectivity of major white matter tracts for drawing learning

Our third analysis tested the selectivity of the tracts identified as the best predictors for drawing learning to drawing. Drawing and visual recognition learning were two different learning outcomes that occurred during the same sensorimotor training session and, therefore, make for a strong test of selectivity to learning outcomes. We hypothesized that the relationship between white matter and individual differences in drawing learning would not transfer to individual differences in visual recognition learning, that tracts predictive of drawing learning would be selective for drawing.

We tested task-selectivity by, first, constructing a new model that we will call the transfer model. The transfer model was a linear model that included only the two predictors selected by the relaxed lasso regression as explaining the largest amount of variance in drawing learning, the left pArc and the left SLF 3, but fit these two predictors to a different dependent measure, visual recognition learning. The original recognition model was a linear model that included only the predictors selected by the relaxed lasso regression as explaining the largest amount of variance for visual recognition learning, including the left MDLFspl and left TPC for the initial dataset and the left MDLFspl for the repeat held-out dataset. Crucially, the transfer and the original recognition learning models both used visual recognition as the dependent variable. We compared the predictive strength of the transfer model to that of the original visual recognition model using a Cox test and a J-test for non-nested models (**Figure 2c**) ^46^. If the transfer model is much worse at predicting visual recognition learning than the original recognition model, such a result would suggest that the drawing learning model does not transfer to visual recognition learning. On the other hand, if the transfer model is just as good or better than the original recognition model, such a result would suggest that the tracts selected as most predictive of drawing learning are not likely specific to drawing.

Results suggest that the relationship between white matter and drawing learning did not transfer to visual recognition learning, and vice versa (**Figure 3c**). The Cox-test demonstrated that the original recognition model (left MDLFspl, left TPC) was a better fit for visual recognition learning than the transfer model (left pArc, left SLF 3), *z* = -3.3122, *p* = 0.0009. Similarly, the transfer model (left pArc, left SLF 3) was not a better fit for visual recognition learning than the original recognition model (left MDLFspl, left TPC), *z* = 0.1382, *p* = 0.8901. Similarly, results in the held-out repeat dataset were similar, *z* = 0.2548, *p* = 0.7988 and *z* = 5.4093, *p* = 6.3×10^−8^, respectively. Results of the J-test were consistent with the results of the Cox test and revealed that combining the transfer and original recognition models to predict visual recognition learning resulted in a model fit that was not significantly better than the original recognition model fit, *t* = 0.227, *p* = 0.821. On the other hand, combining the transfer and original recognition models to predict visual recognition learning resulted in a model fit that was better than the transfer model fit, *t* = 1.686, *p* = 0.099. Results in the held-out repeat dataset were similar, *t* = 0.2169, *p* = 0.8293 and *t* = 1.5153, *p* = 0.1368, respectively. The Cox and J-test results both suggest that models originally selected for drawing learning and visual recognition learning in the RL regression analyses were not transferable to a second learning outcome, indicating selectivity between tracts and learning outcomes.

## Discussion

The current work employed a machine-learning model selection approach to demonstrate selectivity of the mapping between the existing microstructure of major white matter tracts and future learning outcomes. We used diffusion measurements of white matter tissue that occurred over the lifespan to predict individual differences in two learning outcomes that arose from a single sensorimotor training task. The sensorimotor training task consisted of drawing symbols that were previously unknown and the two learning outcomes included: learning to draw the novel symbols and learning to visually recognize those symbols. Results suggest that two left hemisphere white matter tracts, the left pArc and the left SLF 3, selectively predicted individual differences in learning to draw novel symbols but not learning to visually recognize those same symbols. The relationship between the pre-training microstructure of the left pArc and left SLF 3 and drawing learning was found using two independent datasets and two separate statistical analyses (see **Results** and **Supplementary Information**). The relationship between pre-training microstructure and visual recognition learning varied marginally depending on the dataset and statistical analysis but suggested that the pre-training microstructure of the left MDLFspl selectively predicted visual recognition learning. Overall, results suggest that individual differences in the microstructure of human white matter tracts are selectively related to learning outcomes that arise from a single experience.

The current work is the first, to our knowledge, to demonstrate a selective mapping between major white matter tracts and human learning. Tract-selectivity has recently been demonstrated in the murine model ^1–3^, yet evidence for tract-selectivity in humans has remained inconclusive ^4–14^. Furthermore, few studies have tested the mapping between white matter tracts and multiple learning outcomes, leaving the current literature unable to conclude that some tracts are more related to learning than other tasks (tract-selectivity) and more related to learning one task than other tasks (task-selectivity) ^15–22^. Our results add to prior human work that observed widespread changes across multiple white matter tracts during an intensive intervention ^11^. Although interventions often target one learning outcome (e.g., reading), they often impart learning in other domains that are not directly targeted (e.g., attention, social interactions). The onset of an intensive intervention might promote widespread changes ^11^ and our results suggest that only a few of those changes are related to the targeted learning outcome. By using tract microstructure to predict future learning and assessing more than one learning outcome, our results demonstrate a selective mapping between white matter tracts and learning outcomes that likely emerges over time periods much longer than is feasible for interventions.

Two independent analyses demonstrated that the microstructure of two left hemisphere white matter tracts selectively predicted drawing learning: the left pArc and the left SLF 3. The first analysis tested tract-selectivity using a relaxed lasso regression to identify a group of white matter tracts that, together, predicted drawing learning from a set of 22 potential tracts. Each tract entered into the regression was selected based on its unique anatomical connectivity in relation to the functional responses observed during drawing in prior works ^30,34,35,37,38^, including SLF 1 and 2 (combined), SLF 3, pArc, TPC, MDLFspl, MDLFang, Arc, ILF, IFOF, VOF, and FAT in the left and right hemispheres. The relaxed lasso analysis indicated that the left SLF 3 in dorsal cortex and the left pArc in the PVP comprised the group of tracts that explained the most variance in drawing learning (see **Results**). The second analysis tested the task-selectivity of the drawing learning model (i.e., the left pArc and left SLF 3) for drawing by evaluating the model’s prediction strength when predicting a second learning outcome, visual recognition. Results revealed that the drawing learning model did not transfer to visual recognition learning, suggesting a degree of task-selectivity between the microstructure of the left pArc and left SLF 3 and drawing learning. Furthermore, results from both analyses were replicated in a held-out repeat dataset, providing strong evidence that individual differences in the microstructure of the left pArc and left SLF 3 are selectively related to individual differences in learning to draw novel symbols.

Both white matter tracts selected to predict drawing learning were in the left hemisphere, suggesting that learning to draw novel symbols might be supported by left-lateralized communications conveyed along the pArc and SLF 3, consistent with evidence of left-lateralization of functional processes during drawing. Literate adults engage a left-lateralized cortical system during drawing, including regions within the frontal motor, parietal, and ventral temporal lobes ^30,33–35,37–39,47,48^, and these regions are joined by the left pArc and left SLF 3 ^27,29,49^. Furthermore, work in children has demonstrated that the pArc is correlated with individual differences in drawing ability in the left but not the right hemispheres, even after controlling for age ^41^. We and others ^43,50,51^ have suggested that the left-lateralization of white matter supporting drawing learning may be related to the left-lateralization of language processing. In the current study, participants drew symbols that resembled letters of the Roman alphabet and might have relied on the white matter architecture that had been optimized for drawing through extensive life experiences with handwriting letters of the alphabet, an unmistakably language-oriented task. Comparing tract-selectivity in literate and non-literate adults might provide further support for the role of life experiences in adaptive myelination of major white matter tracts that underlie cortical communications during specific learning experiences.

The arcuate fasciculus (Arc) is often segmented into three white matter tracts that allow for at least two different communication pathways, including a long segment (connecting temporal and frontal cortices), an anterior indirect segment (connecting parietal and frontal cortices), and a posterior segment (connecting parietal and temporal cortices) ^27,28^. This segmentation is of great interest to language and reading research because it allows for at least two neural communication pathways during language-oriented tasks: the direct and indirect pathways. The direct pathway is the long segment because it directly connects processing for language perception, such as graphemes (i.e., visual symbols), thought to occur in the temporal cortex with motor processing for language production, such as pronouncing phonemes (i.e., symbol sounds) or writing graphemes (i.e., symbol writing), thought to occur in the frontal motor cortex ^27^. The indirect pathway accomplishes the same connection between perceptual and motor processing but does so indirectly by passing through the parietal cortex. Our results demonstrate that the indirect pathway and not the direct pathway, are predictive of drawing learning. The left pArc and left SLF 3 are effectively the two most posterior segments of the Arc that comprise the indirect pathway ^27–29,52,53^. The SLF 3 is essentially one and the same with the anterior indirect segment of the Arc ^52–54^ and the pArc is essentially one and the same with the posterior segment of the Arc ^29,54^. Our results suggest that parietal involvement might be especially important for drawing learning because we found that the two tracts of the indirect pathway of the Arc selectively predicted drawing learning.

The current work suggests that segmenting the PVP into four white matter tracts may reveal unique relationships with behavior and cortical functioning. The PVP can be segmented into four white matter tracts, however the necessity to segment the PVP into these four white matter tracts has not yet been determined. For example, one study reported no difference in the developmental trajectory of the microstructure of the PVP tracts ^41^. Our results demonstrated that the MDLFspl and pArc selectively predicted different learning outcomes that arose from the same training task. While the MDLFspl predicted visual recognition learning, the pArc predicted drawing learning. Our results are in line with work on the functional mappings in ventral-temporal and parietal cortex. The MDLFspl connects the anterior ventral-temporal cortex with the superior parietal lobe (SPL) where processing is largely associated with visual attention ^55^; the pArc connects the posterior ventral-temporal cortex with the inferior parietal lobe (IPL) where processing is largely associated with visually-guided actions with the hands ^56^. Thus, the MDLFspl may predict visual recognition learning by supporting communication between perceptual processing in anterior ventral-temporal cortex and visual attention in the SPL while the pArc may predict drawing learning by supporting communication between perceptual processing in posterior ventral-temporal cortex and visual processing for hand actions in the IPL. Future work will be necessary to disentangle the mapping between tracts within the PVP and learning.

## Methods

### Participants

Adult participants (18-30 yrs., n = 60) were recruited through flyers posted on the Indiana University campus, online e-flyers, and through word-of-mouth. All participants were screened for neurological trauma, developmental disorders, and MRI contraindications. All participants were right-handed with English as their native language. Participants were compensated with a gift card for each session that they commenced. Data from participants were removed based on signal-to-noise (SNR), motion concerns, or other artifacts (see **Magnetic resonance imaging data analyses**) and, additionally, data from participants whose performance during training and/or testing revealed a lack of engagement were removed (see **Learning rate calculations**), leaving 48 subjects (age: M = 21.21 years, SD = 2.49 years, Range = [18.25, 29.75], 26 F, 22M). All participants provided written informed consent and all procedures were approved by the Indiana University Institutional Review Board.

### Magnetic resonance image acquisition and procedure

Neuroimaging was performed at the Indiana University Imaging Research Facility, housed within the Department of Psychological and Brain Sciences with a 3-Tesla Siemens Prisma whole-body MRI using a 64-channel head coil. Participants were instructed to stay as still as possible during scanning and were allowed to watch a movie or listen to music of their choice during scanning.

T1-weighted anatomical volumes (i.e., t1w) were acquired using a Wave-CAIPI MP-RAGE pulse sequence (TR/TI/TE = 2300/900/3.47 ms, flip angle = 8°, acceleration factor = 3 in phase encoding direction x 3 in slice-selective direction, scan time = 1’14”), resolution = 1 mm isotropic. The T2-weighted anatomical volumes (i.e., t2w) were acquired with a 3D Wave-CAIPI pulse sequence (TR/TI/TE = 2300/900/3.47 ms, flip angle = 8°, acceleration factor = 3 in the phase encoding direction x 3 in slice-selective direction, scan time = 1’15”), resolution = 1 mm isotropic.

Diffusion data were collected using single-shot spin echo simultaneous multi-slice (SMS) EPI (transverse orientation, TE = 87.00 ms, TR = 3470 ms, flip angle = 78 degrees, isotropic 1.5 mm resolution; FOV = LR 210 mm x 192 mm x 138 mm; acquisition matrix MxP = 140 × 128. SMS acceleration factor = 4, interleaved). Diffusion data were collected at two diffusion gradient strengths, with 38 diffusion directions at b = 1,000 s/mm^2^ and 37 directions at b = 2,500 s/mm^2^, as well as 5 images at b = 0 s/mm^2^, once in the AP fold-over direction (i.e., dwi-AP) and once in the PA fold-over direction (i.e., dwi-PA).

Within-session repeat scans were collected for each data type to ensure test-retest repeatability. For each participant, we collected two T1-weighted anatomical images, two T2-weighted anatomical images, two diffusion weighted images with AP phase-encoding, and two diffusion weighted images with PA phase-encoding.

### Magnetic resonance imaging data analyses

All analysis steps were performed using open and reproducible cloud services on the brainlife.io platform ^57^ except for the statistical analyses (see below) that were performed in Matlab R2019b using customized code. All data and analysis services are freely available on brainlife.io (**Table 2**). The code for the relaxed lasso regression is available here: https://github.com/svincibo/learning-white-matter. The code for all other statistical analyses is available here: https://github.com/svincibo/wml-wmpredictslearning.

**Table 2.** Data, description of analyses, and web-links to the open-source code and open cloud services used in the creation of this dataset can be viewed in their entirety here: https://doi.org/10.25663/brainlife.pub.36.

| Application | Github repository | Open Service DOI | Git Branch |
| --- | --- | --- | --- |
| Align T1 to ACPC Plane (HCP-based) | https://github.com/brain-life/app-hcp-acpc-alignment | doi.org/10.25663/bl.app.99 | 1.4 |
| Align T2 to ACPC Plane (HCP-based) | https://github.com/brain-life/app-hcp-acpc-alignment | doi.org/10.25663/brainlife.app.116 | 1.4 |
| Freesurfer Segmentation | https://github.com/brainlife/app-freesurfer | doi.org/10.25663/brainlife.app.462 | 7.1.1 |
| dMRI Preprocessing | https://github.com/brain-life/app-mrtrix3-preproc | doi.org/10.25663/bl.app.68 | 1.7 |
| NODDI model fitting | https://github.com/brainlife/app-noddi-amico | doi.org/10.25663/brainlife.app.365 | 1.3 |
| t1/t2 fitting | https://github.com/brainlife/app-myelin-mapping | doi.org/10.25663/brainlife.app.478 | t1-t2-ratio-v1.0 |
| Denoise mp2rage UNI-T1 | https://github.com/svincibo/app-mp2rage-denoiseUNI | doi.org/10.25663/brainlife.app.506 | main |
| Compute T1 and R1 Maps | https://github.com/svincibo/app-mp2rage-computeT1andR1 | doi.org/10.25663/brainlife.app.514 | main |
| Tractography and DK model fitting | https://github.com/brain-life/app-mrtrix3-act | doi.org/10.25663/brainlife.app.319 | 1.4 |
| Tract Segmentation | https://github.com/brainlife/app-wmaSeg | doi.org/10.25663/brainlife.app.188 | 3.9 |
| Tract Cleaning | https://github.com/brainlife/app-removeTractOutliers | doi.org/10.25663/brainlife.app.195 | 1.3 |
| Tract Analysis Profiles | https://github.com/brain-life/app-tractanalysisprofiles | doi.org/10.25663/brainlife.app.361 | 1.13 |
| Tract Statistics | https://github.com/brainlife/app-tractographyQualityCheck | doi.org/10.25663/brainlife.app.189 | 1.3 |
| Cortex Tissue Mapping | https://github.com/brainlife/app-cortex-tissue-mapping | doi.org/10.25663/brainlife.app.381 | v1.3-snr-input |
| Generate Tract Endpoint Maps | https://github.com/brainlife/app-endpointMapGeneration | doi.org/10.25663/brainlife.app.194 | 1.0 |
| Extract Measures from Tract Endpoints | https://github.com/brainlife/app-cortex-tissue-mapping-stats | doi.org/10.25663/brainlife.app.437 | endpoints-v1.1 |

Anatomical images were aligned to the ACPC plane with an affine transformation using HCP preprocessing pipeline ^58^ as implemented in the Align T1 to ACPC Plane (HCP-based) app on brainlife.io ^59^ for t1w images and as implemented in the Align T2 to ACPC Plane (HCP-based) app on brainlife.io ^59^ for t2w images. ACPC aligned images were then segmented using the Freesurfer 6.0 ^60^ as implemented in the Freesurfer App on brainlife.io ^61^ to generate the cortical volume maps with labeled cortical regions according to the Destrieux 2009 atlas ^62^.

All diffusion preprocessing steps were performed using the recommended MRtrix3 preprocessing steps ^63^ as implemented in the MRtrix3 Preprocess App on brainlife.io ^64^. AP phase-encoded and PA phase-encoded images were combined first and susceptibility- and eddy current-induced distortions as well as inter-volume subject motion were also corrected in this step. PCA denoising and Gibbs deringing procedures were then performed and the volumes were subsequently corrected for bias field and rician noise. Finally, the preprocessed dMRI data and gradients were aligned to each participant’s ACPC-aligned anatomical image using boundary-based registration (BBR) in FSL ^65^.

Diffusion data were removed from the sample if the Signal-to-Noise Ratio (SNR) was less than 15 or if the Framewise Displacement (FD), a widely used measurement of head movement ^66,67^, was greater than 2 mm or if an artifact was apparent. This resulted in a removal of 6 participants.

The microstructural properties of white matter tissue were estimated in a voxel-wise fashion based on preprocessed multi-shell dMRI data. We fit two separate models to the diffusion data: the diffusion kurtosis model (DKI) to the diffusion data to estimate the fractional anisotropy (FA), a summary measure of tissue microstructure that is thought to be related to the integrity of the myelin sheath ^68–70^.

Probabilistic tractography (PT) was used to generate streamlines. We used constrained spherical deconvolution (CSD) to model the diffusion tensor for tracking ^71,72^. Tracking with the CSD model fit was performed probabilistically, using the tractography procedures provided by MRtrix3 Anatomically-constrained Tractography (ACT; ^73–75^ implemented in brainlife.io ^76^. We generated 2 million streamlines at *L*_max_ = 8 and a maximum curvature = 35 degrees, parameters that were optimized for our tractography needs. Streamlines that were shorter than 10 mm or longer than 200 mm were excluded. The tractogram was then segmented using the segmentation approach developed in ^29^ and implemented on brailife.io ^77^. All the files containing the processed data used in this manuscript are available here: https://doi.org/10.25663/brainlife.pub.36.

Streamlines that were more than 4 standard deviations away from the centroid of each tract and/or 4 standard deviations away from the relevant tract’s average streamline length were considered aberrant streamlines and were removed using the Remove Tract Outliers App on brainlife.io ^23,78^.

Tract-profiles were generated for each major tract ^23^ as well as the additional PVP tracts ^29^ using the Tract Analysis Profiles app on brainlife.io ^79^. We first resampled each streamline in a particular tract into 200 equally spaced nodes. At each node, we estimated the location of the tract’s ‘core’ by averaging the x, y, and z coordinates of each streamline at that node. We then estimated FA at each node of the core by averaging across streamlines within that node weighted by the distance of the streamline from the ‘core’. An average white matter measurement was obtained for each tract of interest by averaging across the central 160 nodes, excluding the first and last 20 nodes to avoid partial voluming effects.

### Behavioral procedures

Participants were asked to return for a behavioral session within one week of the neuroimaging session (**Figure 2**). During the behavioral session, participants first performed a 30-minute training session (i.e., Drawing training) followed by a visual recognition test (i.e., Visual recognition testing). An experimenter remained in the room with the participant throughout the behavioral session that was completed within one hour. Code for behavioral procedures can be found here: https://github.com/svincibo/wml-beh.

Stimuli included 200 novel symbols. Using novel, unfamiliar symbols is a well-documented approach that controls for individual differences in pre-training symbol knowledge (James & Atwood, 2008; Kersey & James, 2013; Longcamp et al., 2006, 2008) and allows for a cleaner manipulation of visual, auditory, and motor experience with those symbols. The design and selection criteria for these symbols is described in detail elsewhere (Vinci-Booher, James, & James, 2021). The training required 40 symbols and the visual recognition test required an additional 40 distractor symbols, for a total of 80 symbols. The other 160 symbols were used for counterbalancing; the set of 80 symbols selected for each participant was counterbalanced across participants. Adobe Illustrator was used to create typed versions of these novel symbols. All symbols were in ‘typed’ form in black ink on a white background.

Drawing training: When participants arrived, they were seated at a desk with a digital Wacom writing tablet. Participants were asked to copy novel symbols using the tablet and instructed to make their productions as quickly and as accurately as possible. A Matlab script displayed one of the typed symbols at the top and center of the tablet screen and a box simultaneously appeared below the symbol into which participants were instructed to make their production of the symbol above. Only one symbol was displayed per trial and each trial lasted 4 seconds. Each block included 40 symbols and there were 10 back-to-back blocks, each containing the same 40 symbols. After completing 5 blocks, participants were given a mandatory 3-minute break to rest their hands and eyes before completing the final 5 blocks. The ordering of symbols within each block was randomized. Production duration time was measured for each symbol production trial as the number of seconds between the initial pen-down to the final pen-up.

Visual recognition testing: Participants were asked to perform an old/new recognition test immediately following the training session using an iMac computer and standard keyboard with a key labeled ‘yes’ and a different key labeled ‘no’. Participants first performed a practice session that consisted of individual letters of the alphabet and common shapes (e.g., square, triangle) and the participants were asked to press ‘yes’ for letters and ‘no’ for non-letters. The practice test helped orient participants to the testing context and lasted approximately 2 minutes. After the practice test, participants began the recognition test. During recognition testing, participants were presented with static, typed versions of the 40 learned symbols (i.e., target symbols) along with 40 symbols that were not presented to them during training (i.e., distractor symbols), one at a time and in random order. For each symbol, they were instructed to press ‘yes’ for symbols that they had practiced during training and ‘no’ for non-practiced symbols. Each trial consisted of only one symbol. Each trial began with a 500 ms fixation cross, followed by a 500 ms blank screen, and then a 25 ms stimulus presentation during which a stationary symbol was displayed in the center of the screen. After the stimulus presentation ended, the symbol was replaced by a noise mask until the participant responded or until the trial timed-out. Each trial timed-out after 1 second when participants received feedback that prompted them to respond faster in the next trial (i.e., “Too Slow!”). If the participant responded before the symbol was replaced by the noise mask, the program advanced to the blank screen until the trial time-out criteria was met before moving on to the next trial. Trials that reached the time-out limit were re-presented at the end of the test. Only trials with a participant response (i.e., trials that did not reach the 1-second time-out limit) were used for analyses. Reaction time and accuracy were measured.

### Learning calculations

Drawing learning: Learning rate of the sensorimotor task was calculated by first measuring the amount of time it took a participant to draw a novel symbol, i.e., the draw duration, and plotting this measurement across trials (**Figure 2b**). Trials with a draw duration of 3 standard deviations above or below the within-participant mean were identified as outlier trials and removed. We tested both linear and double exponential models to model the change in draw duration over trials, given that both models have been used in the literature to model learning ^21^. The double exponential models returned fits that were effectively linear despite aggressive efforts at bounding the fits and the linear fits were good fits across participants. The learning rate was calculated for each of the 40 target symbols over the 10 trials as the linear slope of draw duration per symbol over trials. The final learning rate for each participant was calculated by taking the median slope across target symbols for that participant.

Visual recognition learning: Learning to visually recognize each symbol was calculated as the accuracy during visual recognition testing. Learning to visually recognize can be measured by their post-training recognition performance because participants were being tested on novel symbols that they had not been exposed to before they began drawing training. Participants with a visual recognition accuracy of 50% or lower, indicating that they were not performing above chance, were removed, resulting in the removal of 2 participants. We elected to use accuracy and not reaction time to measure visual recognition learning for three reasons: (1) an absence of a speed-accuracy trade-off (b = 0.05, p = 0.57; **Supplementary Information**), (1) an absence of a ceiling effect for accuracy (0.55 < accuracy < 0.91), and (3) slightly greater individual variability captured by accuracy (SD = 0.09) than by reaction time (SD = 0.06).

### Statistical analyses

We were interested in understanding if white matter tracts within the posterior vertical pathway (pArc, TPC, MDLFang, MDLFspl) were more predictive of learning than tracts within the dorsal (SLF3, SLF1and2) and ventral (ILF, IFOF) cortices and, additionally, we were interested in understanding if the tracts that were strong predictors of sensorimotor learning were also predictive of visual perceptual learning. We included three additional control tracts, the vertical-occipital fasciculus (VOF), the frontal aslant tract (FAT), and the arcuate fasciculus (Arc) to control for the fact that the four PVP tracts are vertical tracts while the dorsal and ventral tracts are horizontal. The VOF, FAT, and Arc are vertical tracts that connect ventral cortex with dorsal cortex, but they do not directly connect ventral and parietal cortices. This resulted in a total of 22 tracts of interest, 11 tracts in the left hemisphere and 11 tracts in the right hemisphere.

We used relaxed lasso regression to directly compare among tracts (**Figure 2c**). We entered the average FA for each tract as a predictor in a relaxed lasso model ^31,32^, resulting in 22 potential predictors, one for each tract of interest (see **Methods: *Tract profiles*** for the calculation of average FA of each tract). The lasso procedure selects predictors that, together, explain the most variance in the response variable but is subject to a constraint on the size of the resulting coefficients, effectively shrinking the coefficient estimates. The relaxed lasso removes this shrinkage, de-biasing the coefficient estimates. All variables were standardized by dividing by their own variance to ensure that the magnitude of the beta estimates from the sensorimotor model and the visual recognition model were directly comparable. The best model was selected based on leave-one-out cross-validation. This procedure resulted in two models, one for drawing learning and a second for visual recognition learning.

Additionally, we performed a series of simple linear (marginal) regression analysis to complement the results of the relaxed lasso analysis. Each model included one tract as the predictor and one behavioral measure as the response variable, resulting in 40 simple linear regression models for 20 tracts and 2 behavioral measures (**Supplementary Information**). We tested the significance of the beta-value assigned to the predictor in each model using *t*-test with alpha set to 0.05.

All analyses were applied to the additional repeat diffusion data to support replicability and reproducibility (see **Magnetic resonance image acquisition and procedure**). All statistical analyses were conducted using Matlab v9.11.10 (R2021b), except for the relaxed lasso analysis that was conducted using R v4.2.1 through RStudio 2022.02.1 build 461.

## Funding acknowledgements

This research was supported with grants from the National Science Foundation (OAC-1916518, IIS-1912270, IIS-1636893, BCS-1734853) to Franco Pestilli. Sophia Vinci-Booher was supported by an NSF SBE Postdoctoral Fellowship (SMA-2004877). Daniel McDonald was partially supported by the National Science Foundation (DMS–1753171) and the National Sciences and Engineering Research Council of Canada (RGPIN-2021-02618). Franco Pestilli was also funded by grants from the National Institutes of Health (NIMH R01MH126699, NIBIB R01EB030896, NIBIB R01EB029272), and Wellcome Trust (226486/Z/22/Z), as well as a Microsoft Investigator Fellowship, and a gift from the Kavli Foundation.

## Notes

### Competing Interest Statement

The authors have declared no competing interest.

### Summary of Updates

The revision has been revised to increase the clarity of statements in the Introduction and Results sections.

