## Supplemental Information for "Associative white matter tracts selectively predict sensorimotor learning"

### Supplementary Information

#### Supplemental materials: Results of relaxed lasso analysis using additional AFQ tracts

To ensure that our results were not affected by our hypothesis driven selection of 22 white matter tracts, we conducted our analysis with an extended set of white matter tracts. We included the set of tracts segmented by AFQ [1] with the addition of the four posterior vertical tracts [2] from both hemispheres for a total of 34 tracts. This analysis, therefore, Included the same 22 tracts as in the main document, with the addition of 12 added tracts: uncinate, thalamic radiation, callosum forceps major, callosum forceps minor, cingulum, and corticospinal tract. For drawing learning, results were largely consistent with the findings using the hypothesis-driven entry of 22 tracts into the relaxed lasso regression: the left pArc and left SLF 3 were identified as predictors of drawing learning (**Supplemental Table 1**). However, for visual recognition learning, the relaxed lasso regression failed to identify any tracts that predicted visual recognition learning where the analysis with the hypothesis-driven entry of 22 tracts into the relaxed lasso revealed the left MDLFspl in the original and repeat dataset.

***Supplemental Table 1.*** *Models selected for each learning outcome using relaxed lasso regression.*

| **Response Variable** | **Predictor** | ***β*** | ***S.E.*** | $R^2$ | **Adj.** $R^2$ |
|---|---|---|---|---|---|
| *Drawing learning* | Left pArc | 0.2803 | 0.3219 | 0.0924 | 0.0727 |
| *Drawing learning (repeat dataset)* | Left pArc | 0.2731 | 0.2926 | 0.2210 | 0.1679 |
| | Left SLF3 | 0.2170 | 0.2960 | - | - |
| | Right fronto-thalamic radiation | -0.4252 | 0.3247 | - | - |
| *Visual recognition learning* | - | - | - | - | - |
| Visual recognition learning (repeat dataset) | - | - | - | - | - |

#### Supplemental Materials: Simple linear regression to identify tracts that independently predict drawing and recognition learning

Simple linear (marginal) regression analyses evaluated the relationship between each learning outcome and the microstructure of each tract separately to identify individual tracts (not groups of tracts) that were able to explain a significant amount of variance in learning outcomes. A simple linear regression analysis was conducted for each white matter tract that was included in the relaxed lasso regressions described above and for each learning outcome, resulting in 22 simple linear regressions with drawing learning as the dependent variable and another 22 with visual recognition learning as the dependent variable. Model significance was evaluated using an $F$-test with $p < 0.05$. Results from all significant regressions are reported below and in the **Supplemental Table 2 and** visually displayed with 95% confidence intervals in the **Supplemental Figure 1**. Results from all regressions, including non-significant results, are displayed in the **Supplemental Table 3**.

*Learning to draw novel symbols: left pArc and left SLF3*

The results of the simple linear regression analyses identified the microstructure of only 2 tracts that significantly predicted drawing learning: the left pArc and the left SLF3 (**Supplemental Table 2**). Consistent with the results of the relaxed lasso regression, the relationship between each tract and drawing learning was positive, such that participants with higher FA were participants who were the quickest at learning to draw the novel symbols (**Supplemental Figure 3**), however neither result passed a Bonferonni correction for multiple comparisons, $p < 0.05/22$, i.e., $p < 0.0023$. There were no other tracts that significantly predicted drawing learning, all *p*s > 0.05.

*Visual recognition learning: no significant tracts*

The simple linear regression analyses did not identify any tract that individually predicted visual recognition learning in either the original or repeat data set, all *p*s > 0.05.

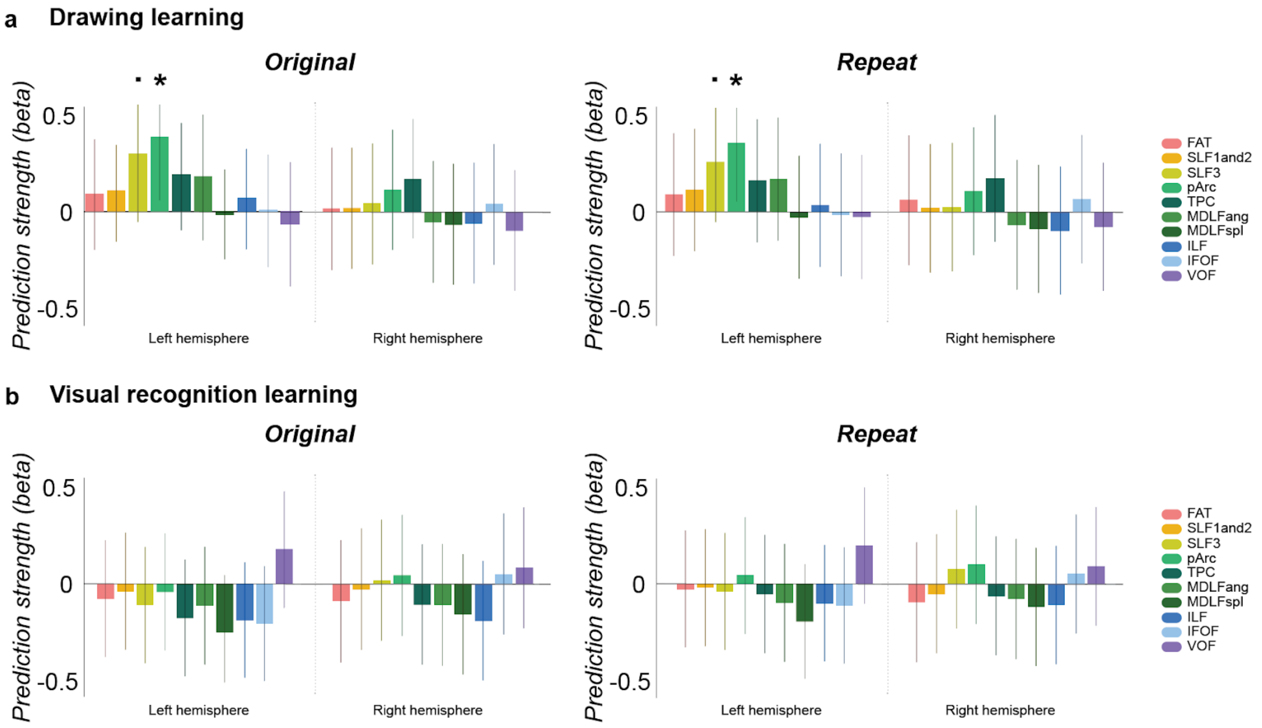


**Supplemental Figure 1.** Simple linear (marginal) regression results: Prediction strength of tract microstructure for drawing and visual recognition learning. **a.** Drawing learning. The left pArc and left SLF3 significantly predicted drawing learning. **b.** Visual recognition learning. There were no tracts that significantly predicted visual recognition learning. Frontal aslant (FAT); superior longitudinal fasciculus, 1st and 2nd segment (SLF1and2); superior longitudinal fasciculus, 3rd segment (SLF3); posterior arcuate fasciculus (pArc); temporal-parietal connection (TPC); middle longitudinal fasciculus connection to the angular gyrus (MDLF-ang); middle longitudinal fasciculus connection to the superior parietal lobe (MDLF-spl); inferior longitudinal fasciculus (ILF); inferior fronto-occipital fasciculus (IFOF); vertical occipital fasciculus (VOF). Error bars represent 95% confidence intervals. ˙ , $p < 0.10$; * , $p < 0.05$.

***Supplemental Table 2.*** *Relationships between one learning outcome and one tract using simple linear regression.*

| Response Variable | Predictor | *β* | *S.E.* | *p* | |
|---|---|---|---|---|---|
| *Drawing learning* | Left pArc | 0.3505 | 0.1381 | 0.0146 | * |
| | Left SLF3 | 0.2745 | 0.1418 | 0.0590 | . |
| *Drawing learning (repeat dataset)* | Left pArc | 0.3308 | 0.1391 | 0.0216 | * |
| | Left SLF3 | 0.2396 | 0.1432 | 0.1010 | . |
| *Visual recognition learning* | - | - | - | - | |
| Visual recognition learning (repeat dataset) | - | - | - | - | |

***Note:*** *All tracts were tested for each learning outcome; however, only significant simple linear regression models are shown here for simplicity. *, $p < 0.05$; . , $p < 0.10$.*

***Supplemental Table 3.***
*Relationships between one learning outcome and one tract using simple linear regression.*

| Response Variable | Predictor | *β* | *S.E.* | *p* | |
|---|---|---|---|---|---|
| *Drawing learning* | | | | | |
| | Left pArc | 0.3505 | 0.1381 | 0.0146 | * |
| | Left SLF3 | 0.2745 | 0.1418 | 0.0590 | . |
| | Left FAT | 0.0827 | 0.1469 | 0.5765 | |
| | Left SLF1and2 | 0.0937 | 0.1468 | 0.5261 | |
| | Left TPC | 0.1701 | 0.1469 | 0.2530 | |
| | Left MDLFang | 0.1617 | 0.1471 | 0.2777 | |
| | Left MDLFspl | -0.0201 | 0.1474 | 0.8922 | |
| | Left Arc | 0.2057 | 0.1443 | 0.1607 | |
| | Left ILF | 0.0593 | 0.1472 | 0.6885 | |
| | Left IFOF | 0.0042 | 0.1474 | 0.9776 | |
| | Left VOF | -0.0567 | 0.1489 | 0.7052 | |
| | Right pArc | 0.1063 | 0.1467 | 0.4720 | |
| | Right SLF3 | 0.0383 | 0.1473 | 0.7957 | |
| | Right FAT | 0.0134 | 0.1491 | 0.9286 | |
| | Right SLF1and2 | 0.0164 | 0.1474 | 0.9119 | |
| | Right TPC | 0.1613 | 0.1455 | 0.2734 | |
| | Right MDLFang | -0.0506 | 0.1489 | 0.7356 | |
| | Right MDLFspl | -0.0624 | 0.1472 | 0.6736 | |
| | Right Arc | 0.1514 | 0.1457 | 0.3043 | |
| | Right ILF | -0.0571 | 0.1472 | 0.6997 | |
| | Right IFOF | 0.0358 | 0.1474 | 0.8091 | |
| | Right VOF | -0.0923 | 0.1468 | 0.5328 | |
| *Drawing learning (repeat dataset)* | | | | | |
| | Left pArc | 0.3308 | 0.1391 | 0.0216 | * |
| | Left SLF3 | 0.2396 | 0.1432 | 0.1010 | . |
| | Left FAT | 0.0841 | 0.1469 | 0.5699 | |
| | Left SLF1and2 | 0.1041 | 0.1466 | 0.4816 | |
| | Left TPC | 0.1506 | 0.1474 | 0.3124 | |
| | Left MDLFang | 0.1574 | 0.1472 | 0.2908 | |
| | Left MDLFspl | -0.0273 | 0.1474 | 0.8541 | |
| | Left Arc | 0.1785 | 0.1451 | 0.2248 | |
| | Left ILF | 0.0310 | 0.1474 | 0.8344 | |
| | Left IFOF | -0.0140 | 0.1474 | 0.9247 | |
| | Left VOF | -0.0247 | 0.1490 | 0.8692 | |
| | Right pArc | 0.0959 | 0.1468 | 0.5166 | |
| | Right SLF3 | 0.0201 | 0.1474 | 0.8919 | |
| | Right FAT | 0.0531 | 0.1489 | 0.7228 | |
| | Right SLF1and2 | 0.0149 | 0.1474 | 0.9199 | |

| | | | | |
|---|---|---|---|---|
| | Right TPC | 0.1551 | 0.1457 | 0.2927 |
| | Right MDLFang | -0.0603 | 0.1488 | 0.6871 |
| | Right MDLFspl | -0.0784 | 0.1470 | 0.5963 |
| | Right Arc | 0.1447 | 0.1459 | 0.3264 |
| | Right ILF | -0.0854 | 0.1469 | 0.5637 |
| | Right IFOF | 0.0579 | 0.1472 | 0.6961 |
| | Right VOF | -0.0688 | 0.1471 | 0.6422 |
| *Visual recognition learning* | | | | |
| | Left pArc | -0.0409 | 0.1473 | 0.7826 |
| | Left SLF3 | -0.1076 | 0.1466 | 0.4666 |
| | Left FAT | -0.0740 | 0.1470 | 0.6171 |
| | Left SLF1and2 | -0.0376 | 0.1473 | 0.8000 |
| | Left TPC | -0.1727 | 0.1468 | 0.2457 |
| | Left MDLFang | -0.1101 | 0.1482 | 0.4613 |
| | Left MDLFspl | -0.2448 | 0.1430 | 0.0936 |
| | Left Arc | -0.0981 | 0.1467 | 0.5069 |
| | Left ILF | -0.1832 | 0.1450 | 0.2126 |
| | Left IFOF | -0.2017 | 0.1444 | 0.1692 |
| | Left VOF | 0.1738 | 0.1468 | 0.2426 |
| | Right pArc | 0.0401 | 0.1473 | 0.7869 |
| | Right SLF3 | -0.0164 | 0.1474 | 0.9121 |
| | Right FAT | -0.0853 | 0.1485 | 0.5687 |
| | Right SLF1and2 | -0.0269 | 0.1474 | 0.8561 |
| | Right TPC | -0.1020 | 0.1467 | 0.4903 |
| | Right MDLFang | -0.1037 | 0.1483 | 0.4880 |
| | Right MDLFspl | -0.1494 | 0.1458 | 0.3107 |
| | Right Arc | -0.1029 | 0.1467 | 0.4863 |
| | Right ILF | -0.1802 | 0.1450 | 0.2204 |
| | Right IFOF | 0.0469 | 0.1473 | 0.7516 |
| | Right VOF | 0.0773 | 0.1470 | 0.6015 |
| *Visual recognition learning (repeat dataset)* | | | | |
| | Left pArc | 0.0424 | 0.1473 | 0.7747 |
| | Left SLF3 | -0.0387 | 0.1473 | 0.7940 |
| | Left FAT | -0.0272 | 0.1474 | 0.8546 |
| | Left SLF1and2 | -0.0196 | 0.1474 | 0.8947 |
| | Left TPC | -0.0527 | 0.1489 | 0.7249 |
| | Left MDLFang | -0.0970 | 0.1484 | 0.5166 |
| | Left MDLFspl | -0.1909 | 0.1447 | 0.1936 |
| | Left Arc | -0.0393 | 0.1473 | 0.7909 |
| | Left ILF | -0.0992 | 0.1467 | 0.5023 |
| | Left IFOF | -0.1091 | 0.1466 | 0.4606 |
| | Left VOF | 0.1938 | 0.1463 | 0.1919 |
| | Right pArc | 0.0944 | 0.1468 | 0.5233 |
| | Right SLF3 | 0.0724 | 0.1471 | 0.6249 |
| | Right FAT | -0.0906 | 0.1485 | 0.5449 |
| | Right SLF1and2 | -0.0501 | 0.1473 | 0.7352 |
| | Right TPC | -0.0595 | 0.6879 | 0.1472 |
| | Right MDLFang | -0.0764 | 0.1486 | 0.6098 |
| | Right MDLFspl | -0.1141 | 0.1465 | 0.4399 |
| | RightArc | -0.0787 | 0.1470 | 0.5948 |
| | Right ILF | -0.1058 | 0.1466 | 0.4741 |
| | Right IFOF | 0.0484 | 0.1473 | 0.1473 |
| | Right VOF | 0.0873 | 0.1469 | 0.5553 |

*NOTE: *, $p < 0.05$; . , $p < 0.10$.*

**Supplemental materials: Participants learned to draw and visually recognize symbols during training**

Drawing is a learning experience that leads to at least two measurable learning outcomes: drawing learning and visual recognition learning. First, drawing practice increases the ability to perform the drawing task itself. As adults practice drawing forms, such as objects, shapes, or symbols, the drawings produced become increasingly recognizable [3]. In young children who are just learning to write letters of the alphabet, writing a letter of the alphabet becomes easier and faster [4,5] and their productions become more legible [6] as they continually practice writing letters. Second, drawing practice leads to changes in the visual processing and memory of the concepts or symbols produced [7–12]. Practice with drawing common objects increased visual recognition of those objects [7] and practice writing pseudo-letters from a novel alphabet increased visual recognition for the practiced pseudo-letters [12]. As an individual practices repeatedly drawing a form, they not only become better at drawing that form but also become better at visually recognizing that form.

To ensure that participants did, in fact, learn to draw and also to visually recognize symbols during the training session, we performed one-sample *t*-test on the sensorimotor learning variable (i.e., slope of draw duration across trials) to confirm that the slope was less than zero (i.e., negative) and also on the visual recognition learning variable (i.e., accuracy) to confirm that it was above chance (i.e., 50%). These analyses confirmed that participants did experience an increase in their ability to draw the symbols throughout the training session and that they also learned to visually recognize the symbols throughout the training session.

***Learning to draw novel symbols***

Participants became faster at drawing symbols throughout the drawing training session, suggesting that participants learned to draw novel symbols during training (**Supplemental Figure 2a**; left). We measured the drawing duration for each symbol drawing trial throughout the 30-minute training session and calculated the slope of draw duration over trials. A one-sample *t*-test on the slope of each participant's drawing durations confirmed that participants' learning slopes ($M$ = -8.6e-4, $SD$ = 1.2e-3) were significantly less than zero, $t(47) = 10.05$, $p$ = 5.06e-4, demonstrating that participants spent less time drawing symbols with each trial that they completed. Additionally, a density histogram of participants' learning slopes demonstrated that the learning slopes for most participants was negative (**Supplemental Figure 2a**; right).

The speed with which participants drew the symbols increased during drawing training, suggesting that participants were learning how to draw the novel symbols during the training session. To our knowledge, learning to draw novel symbols during drawing training has not yet been demonstrated in adult subjects, although it is certainly an intuitive result. One prior work in adults has demonstrated that the drawn productions of common objects became more recognizable with increased practice drawing those objects [3], consistent with the notion that drawing practice improves drawing ability. Additionally, training studies using other sensorimotor learning tasks report similar results: practice with piano playing improves piano playing [13,14] and practice juggling improves juggling [15]. Notably, increases in the speed of drawing occurred without explicit pressure to learn to draw the symbols. We encouraged participants to draw symbols as quickly and as accurately as possible and they were aware that they would be tested on their ability to recognize the symbols after drawing, but they were not aware that we were measuring the duration of their drawings to estimate drawing learning.

#### *Visual recognition learning*

Participants performed the visual recognition test with above-chance accuracy, suggesting that participants learned to visually recognize the symbols during production training (**Supplemental Figure 2b**; left). A one-sample *t*-test confirmed that participants' accuracy during the recognition test ($M$ = 0.76, $SD$ = 0.09) was significantly above chance (i.e., 50%), $t(47)$ = 12.12, $p$ = 4.49e-16, demonstrating that their responses during recognition testing were not likely due to random guessing. Additionally, we visualized reaction times for each participant (**Supplemental Figure 2b**; right) and performed a simple linear regression that revealed no significant relationship between accuracy and reaction time, suggesting the absence of a speed-accuracy trade-off ($R^2$ = 0.048, *beta* = 0.05, $p$ = 0.57).

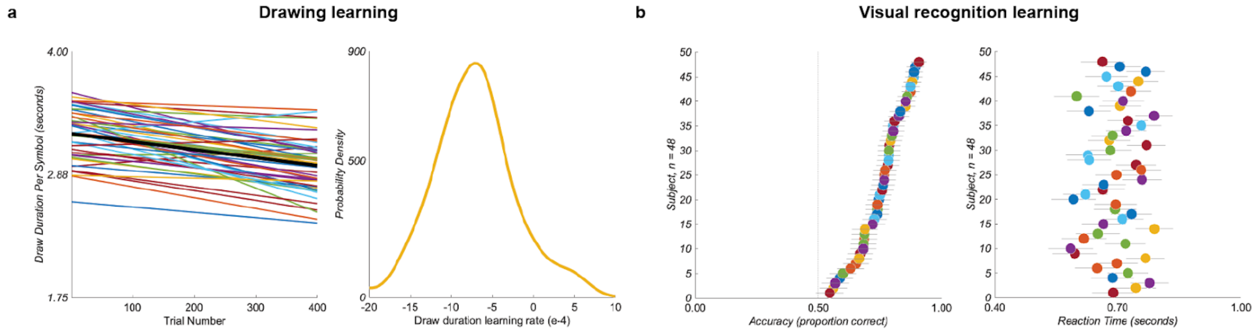


**Supplemental Figure 2.** Behavioral results. **a. Participants learned to draw the symbols.** The draw duration across symbols decreased over the course of the production training (left), $t(47)$ = 10.05, $p$ = 5.06e-4. The white line is the slope of draw duration across trials for all participants. Each non-white line is the linear fit of draw duration across trials for a single participant. The slope of the linear fit of draw duration across trials was negative for nearly all participants (*right*), indicating that the majority of participants learned to draw the novel symbols during training. **b. Participants learned to recognize the symbols.** Participants visually recognized the symbols with above-chance accuracy after production training (*left*), $t(47)$ = 12.12, $p$ = 4.49e-16, suggesting that participants learned to recognize the trained symbols. Each dot represents the proportion correct during the visual recognition test for each individual participant. Average reaction time is also displayed for each individual participant (*right*). Error bars represent standard error across correct trials. For both **a** and **b**, individual subjects are color coded and represented only once per plot.

Prior work has demonstrated that drawing is beneficial for visual recognition learning [7–12], and our results are consistent with the results of these prior studies. In the current study, participants were asked to draw symbols that they had never seen before and were then tested on their ability to visually recognize those symbols. Participants recognized symbols with above chance accuracy after drawing training (**Supplemental Figure 2b**), suggesting that the drawing training contributed to visual recognition learning. However, we did not explicitly manipulate the drawing training and are, therefore, unable to conclude that the drawing training affected visual recognition learning in this study. The drawing training was a copy task in which a typed symbol remained on the screen as a model while participants copied the symbol. It is possible that the above chance recognition accuracy after drawing training resulted from exposure to the model symbol and not the drawing training. Because these were novel symbols, any exposure to the symbols would be expected to lead to above-chance recognition accuracy.

Although it is possible that the above chance recognition accuracy observed after training resulted from only the visual experience of seeing the model symbol, it is highly likely that some of the recognition learning observed after training resulted from drawing. First, prior work has demonstrated that drawing experience facilitates visual recognition of the items that were drawn more than visually perceiving typed symbols [9–12,16,17] and more than other motor activities, such as typing [9,10]. Second, although participants may have learned to visually recognize the symbols from seeing the model symbol, prior research has demonstrated that the act of drawing the symbol has its own effect on visual recognition. In pre-post training studies that have included a model symbol for copying, drawing the symbol beneath a model symbol increased visual recognition more than watching someone else draw the symbol beneath a model symbol, drawing the symbol using a pen without ink beneath a model symbol, watching a symbol unfold on a screen as if being drawn beneath a model symbol, or viewing a static handwritten version of the symbol beneath a model symbol [12,17]. Thus, although we did not include a control condition in this study to determine that the recognition learning was not simply a consequence of exposure to the model symbol during training, prior work using similar study designs suggests that at least some of the recognition learning resulted from drawing.

***No significant relationship between learning to draw and learning to visually recognize***

A simple linear regression was used to determine if the learning rate of symbol drawing was related to visual recognition learning. The predictor was the slope of the draw duration across trials during production training (Figure 4a; left) and the response variable was accuracy (Figure 4b; left). We also tested to see if the slope of draw duration across trials was related to reaction time (Figure 4b; right). We tested the significance of the model using an $F$-test with alpha set to 0.05.

We conducted a simple linear regression analysis to determine if the participants who were quicker at learning to draw symbols were also the participants who were better able to recognize the symbols after drawing. Surprisingly, we observed no significant relationship between learning to draw and learning to visually recognize symbols. Drawing learning was not related to visual recognition learning. A simple linear regression revealed that the learning slope of symbol drawing duration was not a significant predictor of visual recognition learning, as measured by either accuracy ($R^2$ = 0.019, *beta* = 0.05, $p$ = 0.72) or reaction time ($R^2$ = 0.014, *beta* = -0.07, $p$ = 0.55) (**Supplemental Figure 4**). We followed these results with a linear mixed-effects analysis that included random effects for symbol and participant and found similar results. The addition of random effects for symbol and participant significantly improved the model fits; however, the relationship between draw duration slope on visual recognition performance remained non-significant, all $p$s > 0.05.

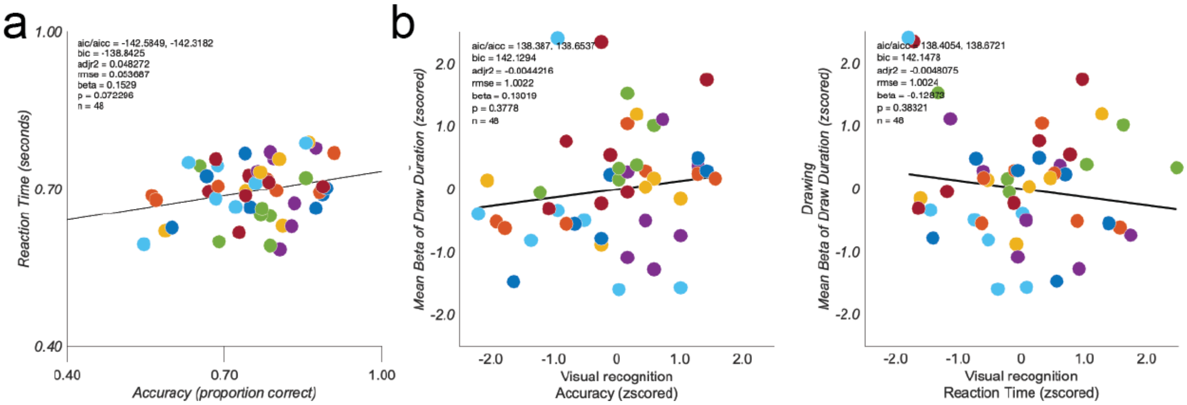


**Supplemental Figure 4.** Relationships among behavioral measurements. **a. Visual recognition learning: no speed-accuracy trade-off.** The relationship between accuracy and reaction time was not significantly different from zero, $b = 0.05$, $p = 0.57$, suggesting that no speed-accuracy trade-off occurred. **b. Non-significant relationship between drawing and recognition learning.** The slope of draw duration over trials was not related to visual recognition accuracy, $b = 0.13$, $p = 0.38$, or reaction time, $b = -0.13$, $p = 0.38$, suggesting that participants who were better at learning to draw the symbols were not the same participants who were better at recognition.
